# Copy number variation contributes to parallel local adaptation in an invasive plant

**DOI:** 10.1101/2024.07.03.601998

**Authors:** Jonathan Wilson, Vanessa C. Bieker, Lotte van Boheemen, Tim Connallon, Michael D. Martin, Paul Battlay, Kathryn A. Hodgins

## Abstract

Adaptation is a critical determinant of the diversification, persistence, and geographic range limits of species. Yet the genetic basis of adaptation is often unknown and potentially underpinned by a wide range of mutational types – from single nucleotide changes to large-scale alterations of chromosome structure. Copy number variation (CNV) is thought to be an important source of adaptive genetic variation, as indicated by decades of candidate gene studies that point to CNVs underlying rapid adaptation to strong selective pressures. Nevertheless, population-genomic studies of CNVs face unique logistical challenges not encountered by other forms of genetic variation. Consequently, few studies have systematically investigated the contributions of CNVs to adaptation at a genome-wide scale. We present a genome-wide analysis of CNV contributing to the adaptation of an invasive weed, *Ambrosia artemisiifolia* – the first such study in an invasive pest. CNVs show clear signatures of parallel local adaptation between North American (native) and European (invaded) ranges, implying widespread reuse of CNVs during adaptation to shared heterogeneous patterns of selection. We used a local principal component analysis to genotype CNV regions in whole-genome sequences of samples collected over the last two centuries. We identified 16 large CNV regions of up to 11.85 megabases in length, six of which show signals of rapid evolutionary change, with pronounced frequency shifts between historic and modern populations. Our results provide compelling genome-wide evidence that copy number variation underlies rapid adaptation over contemporary timescales of natural populations.

**Significance Statement:** Using a population-genomic approach, we identified copy number variants – stretches of DNA that can be either present, absent, or in multiple copies – displaying parallel signatures of local adaptation across the native and introduced ranges of the invasive weed *Ambrosia artemisiifolia*. We further identified 16 large copy number variants, some associated with ecologically important traits including sex allocation and height, that show strong signatures of selection over space, along with dramatic temporal changes over the past several decades. These results highlight the importance of an often-overlooked form of genomic variation in both local adaptation and rapid adaptation of invasive species.

## Introduction

Understanding how populations adapt and persist in the face of rapid environmental change is one of the most pressing challenges of our time. Fundamental to this goal is determining the genetic basis of adaptive evolution. But despite considerable empirical and theoretical work in this area, many questions remain unresolved. For example, does adaptation typically rely on new and beneficial mutations or on standing genetic variation? Does adaptation generally result in the removal or maintenance of genetic variation affecting fitness? Do mutations contributing to adaptation have uniformly small phenotypic effects, or are large-effect mutations important as well? Do populations exposed to similar environments evolve using the same or different genetic variants?

The evolution of quantitative traits was traditionally thought to almost exclusively depend on evolutionary changes at many polymorphic loci with individually small phenotypic effects (1, 2). However, comparatively recent theoretical and empirical studies demonstrate that large-effect variants can also play important roles in adaptation (3–5). Large-effect mutations are particularly likely to contribute to the initial stages of a population’s evolutionary response to a sudden shift in the environment (6), and to facilitate stable adaptive genetic differentiation among populations connected by migration. Such large-effect variants promote local adaptation by resisting the swamping effects of gene flow (7), including cases in which the alleles carry substantial pleiotropic costs (8).

Genomic structural variants, which include inversions, translocations, duplications and deletions, are predicted to have both large phenotypic effects and strong potential to contribute to adaptation (9, 10). Chromosomal inversions have a long history of evolutionary study, initially facilitated by classical cytogenetics (e.g., polytene chromosome studies [(11, 12)]), and more recently through advances in genome sequencing and analysis, which have produced compelling new evidence that inversions often underpin adaptation to environmental change (13–15). These findings have reinvigorated interest in the role of inversions in adaptation, yet other types of structural variation have not garnered the same level of attention.

Several case studies show that copy number variants (CNVs) – structural changes that include gene duplications, deletions and variation in transposable element abundance – have facilitated adaptation in well-characterized systems such as *Drosophila melanogaster* (16), *Anopheles gambiae* (17), and *Arabidopsis thaliana* (18). Prominent examples include the repeated contribution of CNVs to pesticide resistance (19–22), evident in the parallel evolution of CNVs in the agricultural weed *Amaranthus tuberculatus* as a response to glyphosate exposure (23), and amplification of cytochrome P450 family genes in Antarctic killifish and fall armyworm populations exposed to toxins (24, 25). These studies demonstrate the immense adaptive potential of CNVs, yet most are candidate-driven analyses that cannot resolve the broader contributions of CNVs to adaptation. Few studies have systematically characterized the genome-wide contributions of CNVs to adaptive divergence across a range of environmental conditions and stresses (13, 26, 27).

Invasive species have several unique features that make them powerful systems for studying adaptation in nature and uncovering its genetic basis. First, because recently introduced populations are likely to be initially maladapted to local conditions in the new range, there is strong scope for rapid adaptive evolution that is observable within decades (28, 29). Second, some plant invasions have extensive documentation in geo-referenced herbarium collections that can be phenotyped and sequenced to identify and track evolutionary changes over time – an approach rarely possible in natural populations (15, 30, 31). Third, invasive species typically occupy climatically diverse native and invasive ranges, promoting adaptive evolution to local environmental conditions (28, 29, 32). In particular, those with broad ranges further enable tests of the predictability of evolution, since local adaptation across native and invasive ranges may stem from parallel or unique genetic solutions to similar environmental challenges (15, 32, 33). Moreover, they are well suited for evaluating contributions of CNVs, which have been predicted to be important in invasive species adaptation (34, 35). Yet to our knowledge, this hypothesis has never been tested at a genome-wide scale.

Over the last 200 years, the North American native plant *Ambrosia artemisiifolia* (common ragweed) has become a widespread pest on all continents except Antarctica (36). This wind-pollinated, outcrossing species produces highly allergenic pollen that accounts for ∼50% of hay-fever cases in Europe (37). It is also an agricultural pest (38), with glyphosate resistance – a phenotype associated with CNVs in other species (19)– reported in some *A. artemisiifolia* populations (39). Furthermore, climate change is predicted to exacerbate this weed’s impacts, with increased pollen production due to an elongated flowering season (40), as well as reducing the future geographic overlap with key biocontrol agents (41). Previous studies show that *A. artemisiifolia* has established strong signals of local adaptation to climate across its native range, and in introduced ranges in Europe, Asia, and Australia, with rapid local adaptation following each introduction (15, 33, 41–45). We have previously demonstrated a significant contribution of both SNPs and large-effect structural variants – in the form of chromosomal inversions – to climate adaptation in Europe and North America (15). This raises the question of whether other types of genome structural variants (i.e., CNVs) have similarly contributed to adaptive divergence following *A. artemisiifolia*’s range expansion.

In this study, we leveraged a large temporally and spatially resolved dataset to investigate the genome-wide contributions of CNVs to local adaptation in *A. artemisiifolia*. Over 600 whole-genome sequences from individuals collected across the native North American range and the introduced European range, including herbarium samples dating back to 1830, provided a unique opportunity to detect signals of adaptation over space and time. We first investigated whether putative CNVs display signatures of divergent selection in both Europe and North America. By comparing the selection signatures of putative CNVs in each range, we then assessed the degree to which these shared variants evolve in parallel between them. Second, using a subset of 121 phenotyped individuals, we tested for associations between putative CNVs and ecologically important traits, such as flowering time and size. The typically low-coverage and error-prone nature of herbarium sequences renders many existing CNV detection methods unsuitable for these data. As such, we developed a novel approach that combines read-depth and local principal component analysis (PCA) methods to genotype large CNV regions in both modern and low-coverage historic samples, enabling us to identify CNVs, estimate temporal changes in their frequencies, and thus directly track CNV evolution in these populations.

## Results

### CNV identification

To identify CNVs, we calculated read-depth in non-overlapping 10 kbp windows (normalized by the average coverage of each individual sample) across 311 modern re-sequenced genomes spanning both the North American and European ranges of *A. artemisiifolia*. We defined a window as a CNV if at least 5% of samples had average standardized read-depths either greater, or less, than two standard deviations from the window mean across samples. Out of the total 105,175 genomic windows, this resulted in 17,855 candidate CNVs retained for subsequent analyses.

### CNV selection analysis

Local adaptation to spatially heterogeneous environments is expected to leave a signal of extreme allele frequency differentiation among populations (46). As our measures of read-depth for identifying candidate CNV windows were continuous, we used a Q_ST_-F_ST_ outlier test to identify windows with population differentiation in excess of neutral expectations. We first calculated an F_ST_ distribution of neutral SNPs (Figure 1B, D) using the method described by Weir & Cockerham (47). Under neutrality, F_ST_ and Q_ST_ values should have similar distributions, whereas an enrichment of Q_ST_ values within the upper tail of this null F_ST_ distribution provides evidence for local adaptation, with Q_ST_ outliers representing local adaptation candidates (48).

**Figure 1.**
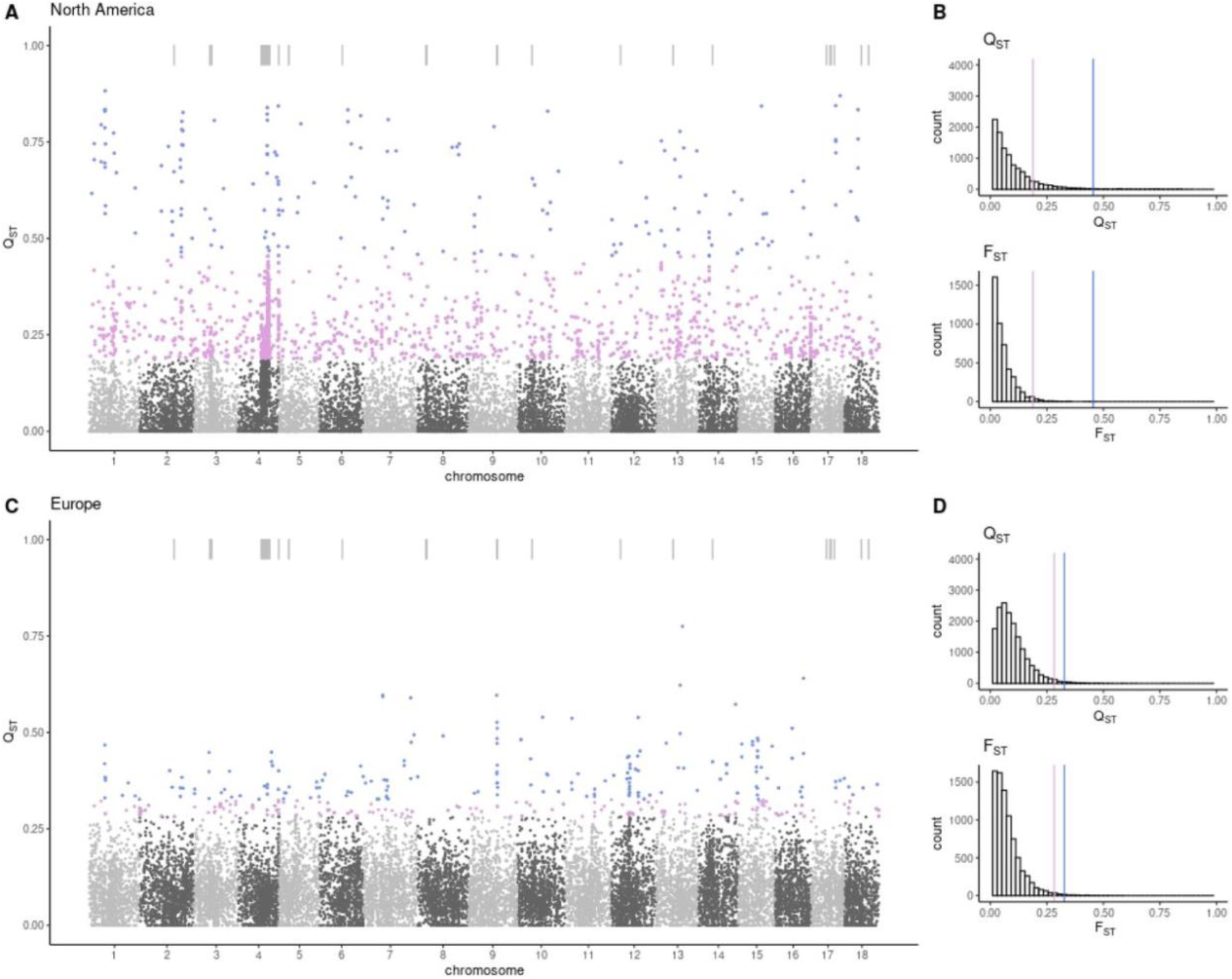
CNV Q_ST_ values indicate regions of selection in the *A. artemisiifolia* genome. A. Q_ST_ values of filtered coverage windows in European populations. Pink windows indicate those above 1% F_ST_ threshold shown in B, blue windows indicate top 1% of Q_ST_ values. C. Q_ST_ values of filtered coverage windows in North American populations. Pink windows indicate those above 1% F_ST_ threshold shown in D, blue windows indicate top 1% of Q_ST_ values. Bars above manhattan plots indicate merged windows > 300 kbp.

In North America, 1,382 CNV windows exhibited Q_ST_ values at or above the top 1% threshold of neutral F_ST_ values: a 7.7-fold enrichment relative to the neutral expectation that 1% of Q_ST_ windows will fall within this tail (p < 2.2e-16; binomial test; Figure 1A, B). In Europe, 339 CNV windows exhibited Q_ST_ values exceeding the top 1% of the F_ST_ distribution: a 1.9-fold enrichment (Figure 1C, D; p < 2.2e-16; binomial test). Using an equation adapted from the McDonald-Kreitman test (49), this excess of outliers is consistent with a true positive rate of 87% for the North American CNV candidate windows, and 47% true positives for European outlier windows (see methods). Of the CNV windows displaying differentiation, 111 were outliers in both ranges (32% of European outliers; p=1.17e-40, hypergeometric test; Figure 2A), a highly significant excess indicative of parallelism in the same CNVs subject to divergent selection in both ranges. In contrast, there was no overlap between the top 1% of neutral SNP F_ST_ values for the two ranges, suggesting that neutral processes cannot explain the parallelism observed in CNVs.

**Figure 2.**
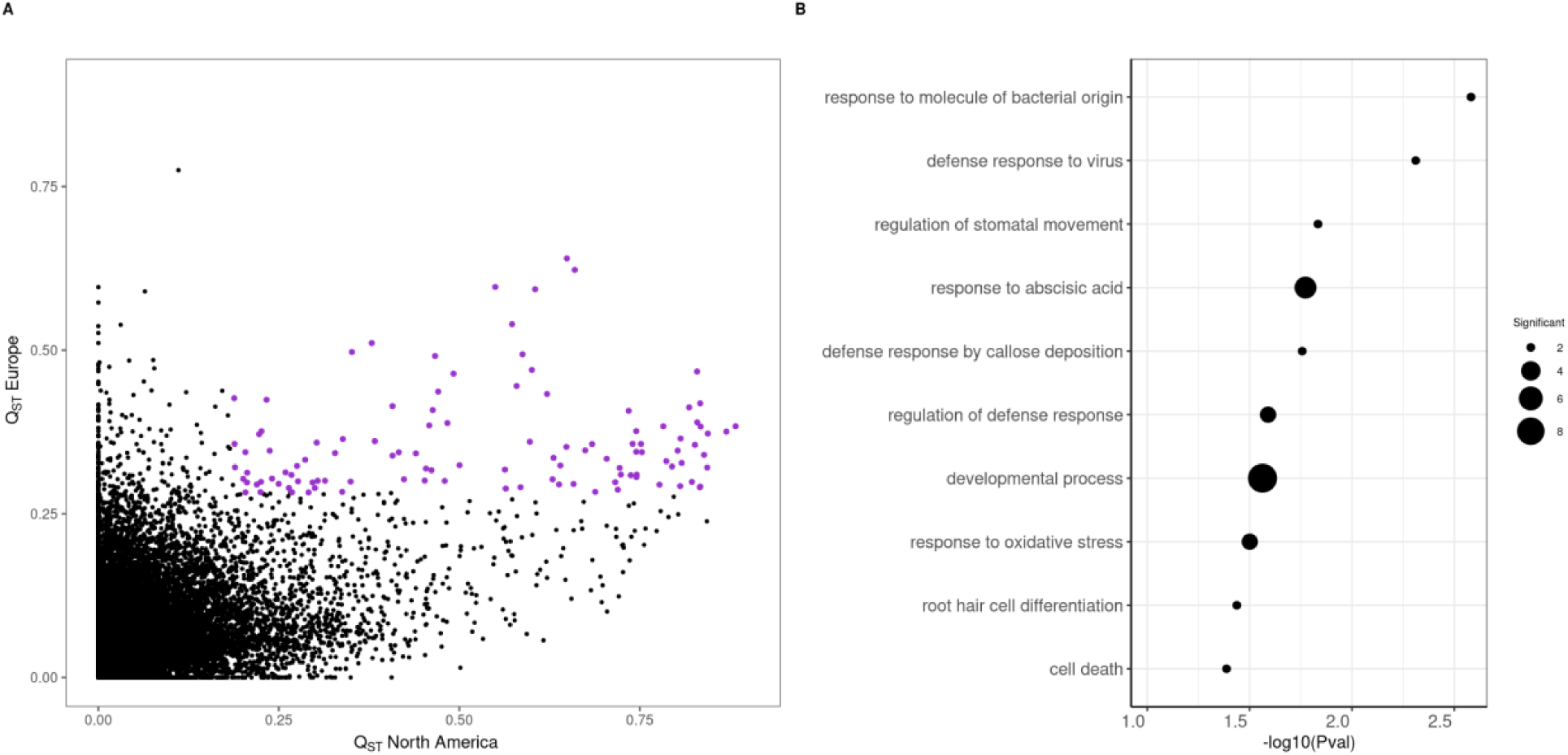
Overlapping Q_ST_ outliers in both ranges and their potential biological functions. A. Distribution of Q_ST_ values of all 10 kbp coverage windows in both North America and Europe. Overlapping windows exceeding the neutral F_ST_ threshold of 1% in each respective range are colored in pink. B. Gene ontology enrichment plot of genes within overlapping outlier Q_ST_ windows, with biological pathways only retained if represented by 2 or more genes (pink in A).

Variation in recombination rate across the genome may interfere with the identification of signatures of selection (50). To account for potential effects of local recombination rate on patterns of CNV differentiation, we separated Q_ST_ windows into three recombination rate bins based on the genetic map described in Prapas et al. (51): low (<0.5 cM/Mbp), medium (0.5-2 cM/Mbp) and high (>2 cM/Mbp). When Q_ST_-F_ST_ analyses were repeated within each recombination rate bin separately, 98.9% and 98.3% of the original Q_ST_ outliers remained significant in North America and Europe, respectively. This demonstrates the minimal impact of recombination rate on the divergent patterns of read depth within CNV windows in this dataset (Supplementary Figure 2). We also investigated the possibility that non-independence between 10 kbp windows drives the observed patterns of divergence and repeatability. To do so, we merged windows with correlated variation in read depth (R^2^ > 0.6) within 1 Mbp of one another. After merging windows, the number of candidate CNVs was reduced from 17,855 to 11,877, with the largest window measuring 11.85 Mbp on chromosome 4. We then repeated the Q_ST_-F_ST_ analysis on these merged windows. Q_ST_ values remained elevated relative to neutral F_ST_ distributions in both North America and Europe (6-fold and 1.3-fold respectively), with 41 outlier windows shared between ranges: far more than expected by chance (hypergeometric test p=2.736e-16).

To identify candidate CNVs associated with climate, we estimated correlations between each candidate CNV window and the six bioclimatic variables (Supplementary Figure 5; 6) that were selected after filtering highly correlated variables from the original 19 WorldClim variables (52). Of the 1,382 significant Q_ST_ windows in North America, 315 (22.7%) were associated with at least one of the six climate variables, whereas only 12 of 339 significant windows in Europe (3%) correlated with climate variables (Supplementary Figures 5; 6), suggesting the primary selective forces driving differentiation of CNVs in Europe are likely not climate-related.

To determine the putative biological functions of the adaptation candidates, we used gene ontology (GO) enrichment analyses of annotated genes residing within the outlier Q_ST_ windows.

Candidate CNVs in North America are enriched for biotic and abiotic stress response genes, with significant GO terms including “systemic acquired resistance,” “response to oomycetes”, and “response to freezing” (Supplementary Figure 3A; Supplementary Table 1). In Europe, significant GO enrichments include the hormonal stress response pathways “response to abscisic acid” and “response to jasmonic acid” (Supplementary Figure 3B; Supplementary Table 2).

Overlapping Q_ST_ candidates between the ranges exhibited GO term-enrichment for the defense response terms “defense response to virus” and “response to abscisic acid” (Figure 2B; Supplementary Table 3).

### CNV-trait associations

We tested for relationships between genome-wide CNVs and 29 ecologically important traits phenotyped in 121 of our samples, each reared in a common garden experiment (phenotype data were previously reported in van Boheemen, Atwater & Hodgins [(43)]). Eighteen traits (Supplementary Table 5) were significantly associated (using a Bonferroni-corrected 0.05 threshold) with normalized read-depth in at least one of the 17,855 candidate CNV windows (Supplementary Figure 4). Of these trait-associated windows, 17 and 4 overlapped with Q_ST_ outlier windows in North America and Europe, respectively. With a more relaxed significance threshold of FDR = 0.05 using the Benjamini-Hochberg method, these overlaps were increased to 76 in North America and 10 in Europe. Of particular interest, two nearby windows on chromosome 14 (h1s14:17180001-17190000, h1s14:18470001-18480000) were associated with flowering onset, dichogamy, and sex allocation (defined as female reproductive biomass/male reproductive biomass; Supplementary Figure 4). These traits display strong latitudinal clines in *A. artemisiifolia*, with overall earlier flowering, much earlier male flowering compared to female flowering, and female-biased sex allocation occurring at high absolute latitudes (43). These two windows flank an annotated gene, AGL-104, that is linked to pollen production in Arabidopsis (53). Moreover, one of these windows (h1s14:18470001-18480000) is a Q_ST_ outlier in North America and Europe, suggestive of parallel divergent selection.

### Large CNV region identification (CNVr)

The continuous measures of read-depth in 10 kbp windows, while reliable in modern samples, were inaccurate when using low-coverage historic data. Since we had previously obtained accurate measures of genotypes for large inversions in these historic samples (15), we implemented a similar approach to identify large CNV regions (CNVr) in order to perform temporal comparisons between the historic and contemporary samples. Furthermore, we would expect many CNVs to be larger than 10 kbp. We therefore used the same linkage disequilibrium-based approach as stated above to identify and merge adjacent windows which appeared to be components of a larger CNV. To corroborate the presence of large segregating CNVs within our dataset, we performed a local PCA of genotype likelihoods in 100 kbp windows across the genome. We determined potential segregating structural variants as regions with at least three adjacent windows that were outliers for distortions in local population structure relative to the rest of that chromosome. This resulted in a minimal size cutoff of 300 kbp for CNVrs. As such, we defined CNV regions as those in which merged read-depth windows (greater than 300 kbp) overlapped with at least three adjacent windows exhibiting variation in local population structure. This approach identified 52 candidate CNVrs.

We then genotyped individuals for CNVrs in our population-genomic data, including low-coverage historic samples. To do this, we used a combination of normalized read depth across the genomic location of each candidate CNVr alongside a PCA calculated within that same region to cluster samples into genotypes differing in both read-depth and PC1 (Figure 3). With this approach, we were able to identify distinct clusters corresponding to genotypes in both modern and historic samples for 16 out of 52 CNVrs. These 16 CNVrs, which ranged in size from 0.3 - 11.85 Mbp, accounted for 8.1% of the 17,885 10 kbp windows identified in the contemporary samples, including 22.6% of outlier Q_ST_ windows in North America and 7.2% of outlier Q_ST_ windows in Europe. Fifteen of these CNVrs exist as heterozygotes within the diploid reference genome, with haplotype 1 containing presence variants, meaning they can be corroborated with alignments between each haplotype of the diploid assembly (Supplementary Figure 9). The high heterozygosity of CNVrs in the reference was likely due to our genotyping method favoring loci with absence alleles that are common in our samples, yet the presence alleles needed to be found in the reference haplotype in order for the CNV to be identified. This is consistent with the low frequencies (mean = 0.196; range: 0.027 −0.479) of all CNVr presence variants (Supplementary Table 6). GO analysis of annotated genes indicates that the 16 CNVrs were enriched for several biological processes, including “methylglyoxal catabolism”, “peroxisome fission”, “pollen tube adhesion”, and “glyphosate metabolism” (Supplementary Table 4).

**Figure 3.**
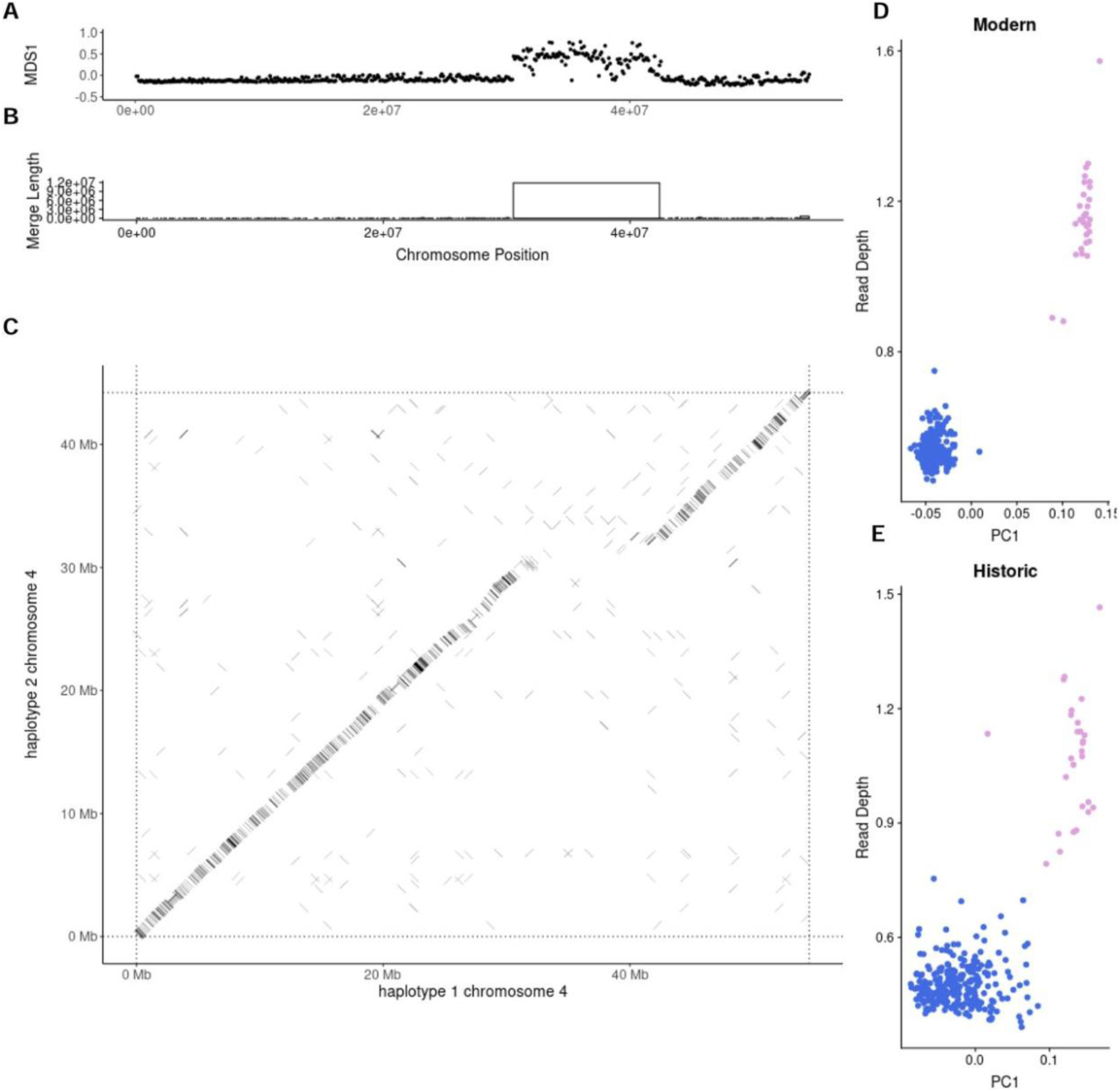
CNVr genotyping of cnv-chr4a. A. MDS coordinates from local PCA in 100kb windows across chromosome 4. Values diverging from zero indicate local structure compared with the rest of the chromosome. B. Distribution of merged window sizes along chromosome 4, with boxes indicating window size on the Y axis and chromosomal position on the X axis. C. Alignment of chromosome 4 Hap1 and Hap2 of the diploid reference genome. A clear gap is present corresponding to local PCA and merged window locations. D, E. Regional PC1 against regional read-depth corresponding to merged window identified in B. Clustering computed using k-means and manual annotation. D. represents modern samples and E. represents historic samples.

The largest CNVr that we identified was cnv-chr4a, which we estimated to be 11.85 Mbp in length. Closer inspection of this region within the diploid reference reveals that large regions on haplotype 1 are absent on haplotype 2, but are interspersed by three smaller inversions (Figure 3; Supplementary Figure 8; 9). Analysis of coverage depth across chromosome 4 of three closely related Ambrosia species sequences mapped to the *A. artemisiifolia* reference revealed the absence haplotype as the likely ancestral state (Supplementary Figure 10). The derived insertion variant contains an excess of transposable elements (TEs) relative to the remainder of chromosome 4 (87.21% versus 70.48%). TE family Ty1/Copia accounts for 32.45% of the TEs within cnv-chr4a, compared to just 14.87% throughout the rest of this chromosome. Repetitive elements display greater density towards the beginning of this region (Supplementary Figure 11), where gene density is very low. The region toward the end of the cnv-chr4a, which exhibits greater gene density, shows strong synteny with chromosome 2 and aligns with inversions present in the reference alignment (Supplementary Figure 8; 11). Most of chromosome 4 displays synteny with chromosome 2, suggesting they are homoeologous chromosomes. The largest gap in syntenic blocks corresponds to the gene-depleted and TY1/Copia-enriched region of cnv-chr4a (Supplementary Figure 11). It is therefore likely that this complex structural variant consists of a series of inversions which have subsequently been separated by a large TE expansion. Recombination is likely strongly suppressed within this structural variant, as the coverage windows exhibit strong linkage disequilibrium across the region. Candidate CNV windows within cnv-chr4a are Q_ST_ outliers (Figure 1A; C) and associated with the bioclimatic variable mean diurnal range (Supplementary Figure 5), consistent with the CNVr contributing to local adaptation.

Another noteworthy CNVr, cnv-chr8a, contains an ortholog of the *Arabidopsis thaliana* EPSPS locus. This CNVr lies within a large inversion, hb-chr8, previously described in Battlay et al. (15). While the frequency of cnv-chr8a is not strongly correlated with the frequency of this inversion (R^2^=0.006), cnv-chr8a presence alleles occur exclusively on the common, and presumably ancestral, orientation of the inverted region.

### Spatio-temporal CNVr modeling

We used whole genome sequences derived from >284 historical herbarium samples (dating back to 1830) and generalized linear models (GLMs) to uncover how the 16 CNVr alleles may have changed in frequency over both space and time. These GLMs predicted genotype as a function of range (native or introduced), latitude and year of collection, and each model was reduced to remove any non-significant interactions between these variables. To correct for population structure, we also included the first principal component of genetic variation (calculated from 10,000 neutral SNPs) as a covariate in each model. Twelve of the 16 CNVrs displayed at least one significant predictor variable (time, range, latitude or interactions). Nine CNVrs exhibited significant associations with latitude (cnv-chr4a, cnv-chr4b, cnv-chr5a, cnv-chr9a, cnv-chr14a, cnv-chr17b, cnv-chr17c and cnv-chr18a) (Supplementary Table 7; Supplementary Figure 12).

Models of three CNVrs (cnv-chr10a, cnv-chr14a and cnv-chr17a) contained significant three-way interactions (Figure 4; Supplementary Table 7; 8; Supplementary Figure 12). Of note, both cnv-chr14a and cnv-chr17a displayed clinal patterns in North America regardless of year, whereas this same latitudinal pattern was present only in modern European samples — strong evidence of clinal reformation following an initial period of maladaptation in the introduced range (Figure 4; Supplementary Table 7; 8). The large cnv-chr4a insertion displays latitudinal clines in modern populations across both ranges, with the insertion at higher frequencies at lower latitudes. In the native range, this appears to be driven by increasing frequencies over time in more central and southern populations of North America (Supplementary Table 8).

**Figure 4.**
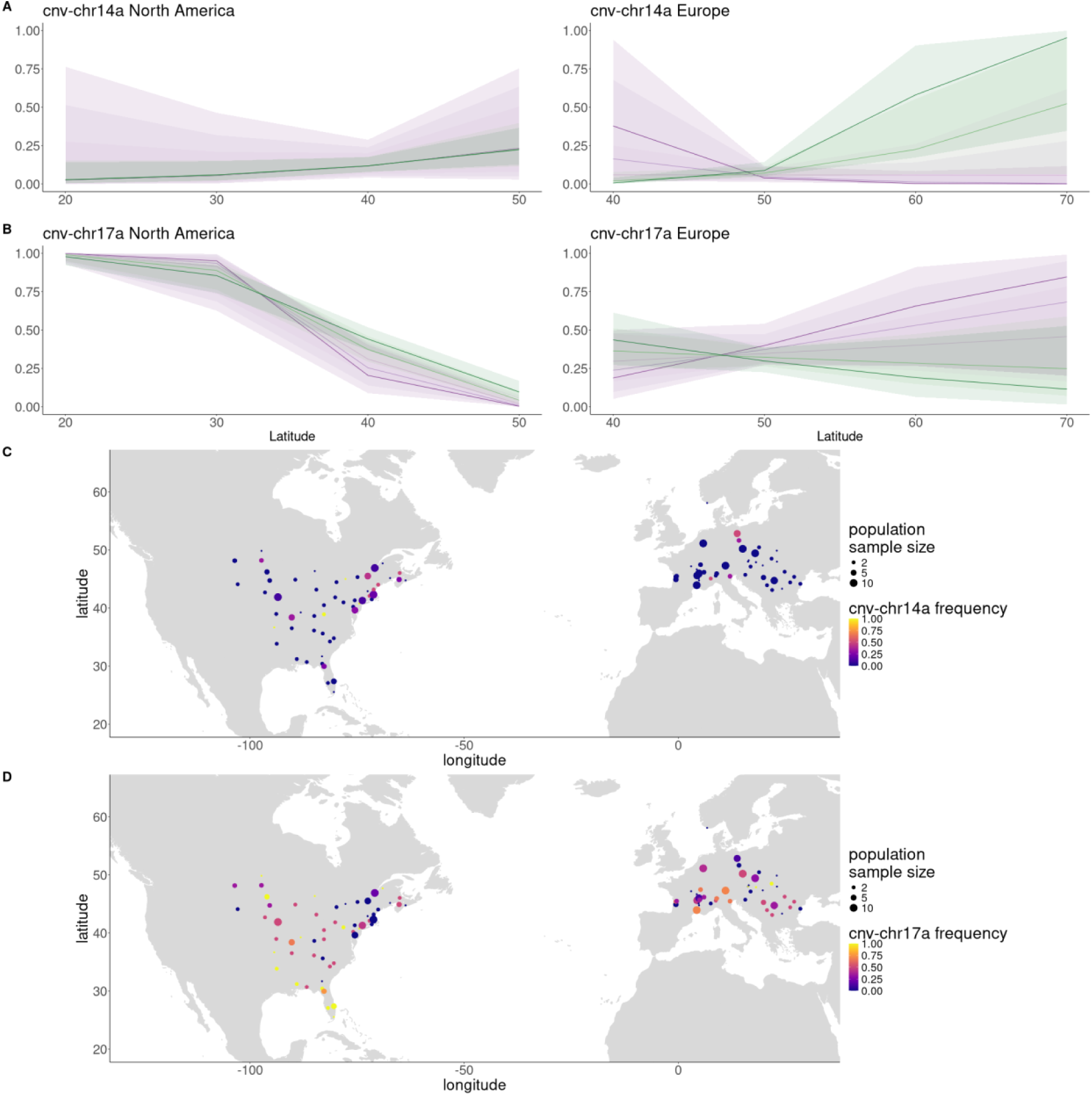
Spatio-temporal CNV frequency shifts. A, B. Logistic regression models for A) cnv-chr14a and B) cnv- chr-17a with error bands representing 95% CI of least-squares regressions of CNV frequency. Time binned into five categories, ranging from historic (purple) to modern (green). Model information found in Supplementary Table 6. C, D. Frequency of C) cnv-chr14a and D) cnv-chr17a in modern *A. artemisiifolia* populations.

Correspondingly, this CNV overlaps with a previous SNP-based result consistent with a selective sweep in the St. Louis population (15).

To further analyze the potential adaptive significance of these CNVrs, we assessed associations between each variant and the same 29 phenotypes analyzed above (43). The cnv-chr10a variant displayed Bonferroni-significant associations with total biomass, root weight and total number of male inflorescences, whilst cnv-chr14a was significantly associated with sex allocation (Supplementary Table 9).

## Discussion

CNVs are increasingly recognized as important in local adaptation (13, 24, 26, 27, 54). However, previous empirical evidence is predominantly limited to examples of pesticide resistance (18, 25) and candidate gene studies (55, 56). Genome-wide analyses of CNVs at the population scale are rare beyond model organisms (54). While many methods exist for identifying CNVs from resequenced genomes (57, 58), these often rely on long reads, or short reads with higher coverage than available for many population genomic datasets, including our historic specimens. We therefore combined local read-depth, linkage disequilibrium, and deviations in population structure along the chromosome, to identify CNVs in modest-coverage samples collected over the past 190 years.

We implemented a genome-wide discovery approach to identify CNVs and examined their potential contributions to local adaptation across the native and an invaded range of *A. artemisiifolia*. CNV windows were enriched for signatures of local adaptation, which occurred disproportionately in parallel between native and invasive ranges. As this signal was not replicated in putatively neutral SNP loci, this implies CNV-driven local adaptation across Europe takes place via standing variation inherited from North America. Furthermore, large CNV regions identified in modern and historical samples, such as cnv-chr14a and cnv-chr17a (Figure 4), show evidence of rapid local adaptation across the short timescale (ca. 150 years) of *A. artemisiifolia*’s invasion in Europe.

*Ambrosia artemisiifolia* CNV windows exhibited extensive signals of local adaptation, including elevated geographic divergence in fold coverage, relative to neutral expectations, in both North America and Europe (Figure 1). These windows contain an over-representation of genes involved in abiotic and biotic stress responses in North America (Supplementary Figure 3A, Supplementary Table 4) and pathogen defense in Europe (Supplementary Figure 3B, Supplementary Table 3) – consistent with SNP-based F_ST_ outlier windows in Europe being enriched for defense related functions (30). The overall weaker patterns of CNV differentiation in Europe relative to North America are also consistent with previous SNP-based analyses showing fewer differentiated outlier SNP windows in Europe (15). Some CNVs were associated with traits important to local adaptation, including a CNV on chromosome 16 associated with mature plant height (Supplementary Figure 4), which also overlaps an annotated NB-ARC domain. NB-ARC domains, occurring in most plant resistance (R) genes, are involved in nucleotide binding and recognition (59), and duplications of such genes underlie the evolution of resistance to pathogens (60, 61). CNVs also appear to influence phenological traits. For example, the CNV windows on chromosome 14 are associated with flowering time (Supplementary Figure 4). Overall, our data point to important roles of CNVs in the local adaptation of *A. artemisiifolia*, which aligns with evidence from other species that structural variants widely contribute to adaptation (62–64).

One third of adaptive CNV windows in Europe were also candidates for local adaptation across North America. Previous work in *A. artemisiifolia* has revealed similar patterns of repeatability, or parallelism, with respect to SNPs (15, 33), inversions (15), and genes affecting locally adapted traits like flowering time and sex allocation (43). Invasive species are expected to evolve in parallel when responding to analogous selection pressures, as observed in our system and in others, such as *Drosophila suzukii* and European starling (65, 66). Such parallelism is promoted when standing genetic variation involved in local adaptation in the native range is recruited as a source of adaptive variation within the invasive range (67, 68). Multiple introductions from several genetically diverse source populations from North America to Europe presumably facilitated repeatability by ensuring that most of the important standing variants successfully made the journey to the new range (30, 33). That all CNVrs were present in historical European populations indicates that they were introduced into Europe during the early stages of invasion (Supplementary Table 6).

The 16 large CNV regions that we detected using a combination of read-depth, linkage disequilibrium and local population structure analyses (Figure 3A, Figure 3B) contained 22.6% of Q_ST_ outlier windows from North America and 7.2% of the outlier windows from Europe. Yet these CNVrs comprised only 0.23% of the genome, demonstrating their disproportionate contribution to these signals of local adaptation. Remarkably, 15 of the CNVrs co-localize with segregating presence/absence variants in the highly heterozygous diploid reference, supporting our detection method. Closer analysis of the largest CNVr (cnv-chr4a) within the reference reveals that our detection method may lack sensitivity in fully revealing structural complexity within CNVrs (Figure 3). Multiple abutting inversions and CNVs within this region appear to segregate together as a single, complex structural variant. Nevertheless, the regions we identified demonstrate the general adaptive potential of structural variants, in which CNVs are predominant features. Since our method of detection for CNVrs was biased towards identifying large CNVrs with presence variants on haplotype 1 of the reference, and our genotyping method was biased towards identifying loci with common absence alleles, our results represent a lower bound on the prevalence and adaptive significance of CNVrs in *A. artemisiifolia*. Investigations using pangenomics (69) to elucidate a more complete picture of the contribution of CNVs to adaptation are therefore warranted.

Our study goes beyond the traditional population-genomic approach of detecting signals of local adaptation using contemporary samples alone. Our use of historical sequence data also allowed us to track temporal change in CNV frequencies across nearly two centuries – a period that spans the establishment and spread of *A. artemisiifolia* within Europe (36, 70), and significant environmental upheaval in both ranges, owing to industrialization, agriculture, and climate change (71). CNVs are rarely genotyped in historic genomes because sequence quality is poor (72). However, by focussing only on large CNVs identified using modern data we were able to confidently assign genotypes in historic samples. This use of modern data to validate historic sequence variant calls is common in temporal genomic studies (30, 73). Eight CNVrs display clear frequency shifts over time, consistent with rapid adaptation over its recent evolutionary history. For example, while cnv-chr14a and cnv-chr17a both exhibit a consistent latitudinal cline in historical and modern North American populations, these clines are only evident in modern European populations (Figure 4A; Supplementary Table 7; 8), which is consistent with the rapid cline reformation in Europe following an initial period of post-introduction maladaptation. The cnv-chr14a CNVr is associated with sex allocation (Supplementary Table 9), and candidate CNV windows within the region exhibit associations with flowering time, sex allocation and dichogamy (Supplementary Figure 4), traits which show parallel clines in Europe and North America (43). Furthermore, cnv-chr14a lies within 20 kbp of the AGL-104 gene, which is involved in pollen development in Arabidopsis (53). Flowering time is a complex trait that is affected by diverse forms of genomic variation (74) – our results indicate that SNPs, inversions (15), and CNVs each play important roles in the rapid adaptation of this important trait in *A. artemisiifolia* populations.

Many well-characterized CNVs in other species are associated with the evolution of pesticide resistance (19, 20, 25). We identified a CNVr (cnv-chr8a) potentially involved in herbicide resistance. An ortholog of the *Arabidopsis thaliana* EPSPS locus, the molecular target of glyphosate herbicides, lies within the cnv-chr8a region. CNVs confer resistance to glyphosate in numerous other weed species, where the increased gene expression caused by EPSPS gene amplification ameliorates the herbicide’s toxic effects (19, 55). Glyphosate resistance has been documented in some *A. artemisiifolia* populations (39), and while we do not know which populations in our study might be glyphosate resistant, this CNVr is a strong candidate for future study.

Previous population-genomic analyses of *A. artemisiifolia* provide strong evidence that SNPs and putative chromosomal inversions contribute to local adaptation; here we provide evidence of a similar role for CNVs. Although CNVs are known to have large effects on traits (17), we cannot be sure that the variants we have identified are the direct targets of selection – they may instead be in linkage disequilibrium with other variants that are the actual targets. Assessing relationships between CNVs and nearby SNPs is fraught, because CNVs disrupt SNP calling (75). Two CNVrs overlap inversions identified in Battlay et al. (15), but are not in strong LD with the inversion genotypes (R^2^ = 0.006-0.01). However, smaller CNVs may exhibit stronger associations with other SVs as part of coadapted gene complexes (76) or neutral hitchhikers. Large insertion-deletion variants result in may hemizygous regions of reduced recombination which may collect and bind together multiple variants (77). Functional assays such as RNA-seq experiments are required to understand the mechanistic effect of these CNVs on traits and fitness (78). Further efforts to determine the functional effects of CNVs, alongside greater sample sizes, would also help uncover the likely epistatic interactions between CNVs and other adaptive variants. The potential existence and nature of these interactions are pertinent questions in evolutionary biology lacking empirical investigations on genome-wide scales (79). Future work should also consider the role of other forms of genomic variation, for example transposable element abundance and genome size which have been linked with local adaptation and aggressive range expansion (74), alongside investigating the roles of CNVs in biotic interactions, such as pathogen response.

Our study highlights the importance of copy number variation in the evolution of a widely distributed and rapidly adapting invasive weed. While CNVs have previously been implicated in adaptation in response to specific selection pressures in other species (23, 24), our genome-wide discovery approach was able to identify candidate genomic regions that are more broadly representative of the contribution of CNVs to local adaptation. We have linked several of these candidates with traits ranging from flowering time to pathogen resistance. Along with previous studies showing that SNPs and chromosomal inversions underlie local adaptation during *A. artemissifolia*’s expansion across vast environmental gradients, these new findings make it clear that CNVs account for a significant and previously unrecognized component of this plant’s past success and are consequential for its invasive capacity wherever it may be introduced in the future.

## Methods

### Samples and alignments

Analyses were conducted on 613 whole-genome *Ambrosia artemisiifolia* sequences described in Bieker et al. (30) and a chromosome-level, phased, diploid *Ambrosia artemisiifolia* genome assembly (15). Alignments and SNP calls against the primary haplotype of the diploid reference (haplotype 1) were generated by Battlay et al. (15), using the Paleomix v1.2.13.4 (80) pipeline and GATK UnifiedGenotyper v3.7 (81). Modern and historic samples were sequenced from across the species’ native North American (155 modern and 92 historic samples) and introduced European (156 modern and 192 historic samples) ranges. Modern samples were collected between 2014 and 2018 and sequenced to a mean coverage of 2.9x. Historic samples were sequenced from herbarium samples collected between 1830 and 1973 with a median collection date of 1905 (Supplementary Figure 1), and sequenced to a mean coverage of 1.4x. Present-day population samples, whose geographic coordinates were recorded during sampling, included between n = 1 to n = 10 individuals, however populations where n = 1 were removed from analyses requiring population level information. Historic individuals were grouped into populations according to their proximity (15, 30). Cases where only one sample was obtained from a geographic location were excluded from analyses in which population information was a requirement. Additionally, we aligned sequences of three other Ambrosia species (30) to the primary haplotype of the diploid reference using the Paleomix pipeline, as described in Battlay et al. (15).

### Depth of coverage analysis

In order to identify copy number variation within our resequenced common ragweed individuals, we analyzed depth of coverage in non-overlapping 10 kbp windows using Samtools v1.9 depth (82) on alignment bam files. In the initial analyses of modern samples we only used reads with mapping quality > 30 (-Q 30). Subsequent read-depth analyses of historic samples used reads with mapping quality > 5 (-Q 5) in order to accommodate their poorer mapping quality. Each window was then normalized by dividing window depth by the genome-wide coverage for the sample. To apply a population frequency-based filter to this dataset, we kept only windows which had at least 5% of samples greater than or less than 2 standard deviations from the population mean. This filtering procedure resulted in 17,855 of 105,175 windows (5.8%) being classified as copy number variants (CNVs).

### Q_ST_-F_ST_ analysis

Genomic loci that have responded to spatially heterogeneous selection are expected to show elevated differentiation among populations, relative to neutrally evolving loci. SNPs associated with local adaptation can be detected as outliers of genome-wide scans of F_ST_ or similar statistics (47, 83, 84). However, unlike SNP data, candidate CNVs have been identified by depth of coverage represented on a continuous scale. As such, coverage can be viewed as a phenotypic measurement that is analogous to measurements of continuous traits. Tests for trait responses to divergent selection are often inferred using a Q_ST_-F_ST_ analysis (85, 86). Theory predicts that Q_ST_ values for neutrally evolving traits should follow the same distribution as F_ST_ for neutrally evolving loci (48). We used coverage data to measure Q_ST_ values for each window (analyses were conducted separately for the European and North American ranges), adapting the relationship between Q_ST_ and population variation from Whitlock (48):

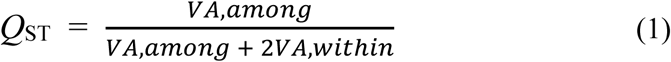

where VA,among represents the among-population variation in coverage for a given window, and VA,within represents within-population coverage variation.

We performed linear mixed models, using the lme4 package in R (87) on populations from each range to extract variance components attributed to within- and among-population variation for each coverage window. Unlike other analyses performed in this study, we used unnormalised coverage for each window in the model, and accounted for variation in individual sample depth by including individual median coverage as a covariate in the model, and population was included as a random effect. The variance among populations (VA,among) was extracted as the variance component attributed to the population, and within-population variance (VA,within) corresponded to the model’s residual variance component.

To identify Q_ST_ values with divergence in excess of neutral expectations, the distribution of Q_ST_ values in each range was compared to the distribution of neutral F_ST_ values from the same populations. F_ST_ distributions were calculated in VCFtools (88) using 10,000 putatively neutral and independently segregating LD-pruned SNPs, randomly sampled from outside both genic regions and known structural variants (15). Outliers were classified as windows with Q_ST_ values greater than 99% of neutral F_ST_ values. Under neutrality, 1% of Q_ST_ values are expected to fall above this 99% threshold, and we therefore tested whether there was a significant excess of windows above this null expectation. To identify the rate of true positives, we used the following calculation

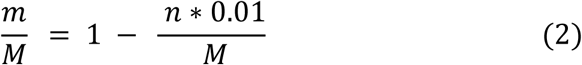

where *n* represents the total number of windows analysed, *M* represents the total number of outliers, *m* represents the number of true positives, and therefore *M* - *m* represents the number of false positives. Analyses were performed using modern data from North America and European ranges separately. We used a hypergeometric test to identify significant overlaps in both Q_ST_ and F_ST_ loci between both ranges.

### Environmental-CNV Associations

Correlations between genetic variation and environment provide further evidence for local adaptation on the genomic level. CNV-environment associations were identified by measuring Spearman’s correlation coefficients (Rho) between normalized coverage of each 10 kbp window and climate variables. Climatic variables were extracted from WorldClim (52) for each of the geographic coordinates of our sample of individuals. We excluded highly correlated variables (R^2^ > 0.7) from the analysis, resulting in six variables: BIO1: annual mean temperature, BIO2: mean diurnal range, BIO8: mean temperature of wettest quarter, BIO9: mean temperature of driest quarter, BIO12: annual precipitation, and BIO15: precipitation seasonality. Outliers were identified as the top 1% of the empirical p-value of the Rho distribution. Coverage windows that were in the 1% tails of the CNV-environment distribution and outliers in our Q_ST_ analysis were considered CNV-climate adaptation candidates.

### Identifying large CNV regions

Since larger CNVs will affect coverage in multiple adjacent windows, we used a linkage disequilibrium-based approach to merge nearby windows. We merged windows within 1 Mbp (and all windows within this region) that had sample depths that were correlated at R^2^ > 0.6. By generating heatmaps of LD between CNV windows, we visualized the presence of correlated read-depth, indicative of larger CNVs (Supplementary Figure 7). This resulted in 11,877 windows ranging in size from 10 kbp to 11.85 Mbp.

Local principal component analysis has previously been used to detect population-genomic signatures of chromosomal inversions (15, 62, 89). Here we employed this method to identify distortions in local relatedness due to CNVs using two modifications. Due to low-coverage and bias in historic samples, we performed these steps initially on modern samples only, including historic samples for genotyping later (see below). Firstly, we calculated local covariance matrices for each window using ANGSD (90) and PCAngsd (91). Secondly, we did not include a filter for missing data in our ANGSD command, which meant that missingness would cause distortion in the local PCAs. The analysis was performed in non-overlapping windows of 100 kbp, and multidimensional scaling axes 1-4 (calculated from local PCAs across each chromosome) were examined for blocks of outliers indicative of large structural variants. While individual MDS outlier windows are not necessarily due to structural variations, SVs are the most likely cause of signals that are consistent across adjacent windows. Therefore, we retained candidates that included at least three adjacent windows that were outliers, so that the lower limit of CNVr size was 300 kbp. As such, we identified candidate CNVr as those where MDS candidates overlapped the merged read-depth-based CNV windows which were also greater than 300 kbp in length. These 52 candidate CNVrs corresponded to 68% of the merged CNV windows greater than 300 kbp in length.

### CNV region genotyping

We attempted to genotype modern and historical samples, separately, for each of the 52 candidate CNVr. To do so, we performed another local PCA across the length of each candidate CNVr using PCAngsd (91). We then compared PC1 of the candidate CNVr against the average normalized read-depth across that CNVr to identify whether individuals clustered by read-depth and PC1. To genotype these samples according to these clusters, we used k-means clustering from the ClusterR package in R (92), followed by manual annotation of genotypes. Samples appeared to either cluster into groups of two or three genotypes. CNVr candidates displaying two clear genotypes were likely to have one rare homozygote, or the heterozygote and one homozygote class were indistinguishable. As such, we assigned k-values of either 2 or 3 depending on whether visual inspection indicated the presence of two or three segregating genotypes. To test for association between CNVrs overlapping or neighboring chromosomal inversions, we used the cor function in R (R team) to calculate correlations between CNVr and the genotypes of overlapping inversions that were previously identified by Battlay et al. (15). To visualize CNVrs that were heterozygous between haplotypes of the diploid reference genome, we aligned both reference haplotypes using minimap2 v2.1.8 (-k19 -w19 -m200) (93) and generated dotplots of the alignments. To call segregating SVs in the diploid reference, we aligned both haplotypes using nucmer (-- maxmatch -c 100 -b 500 -l 50) within the mummer v3.23 software package (94). We then used SyRI (95) to identify and plotsr (-s 300000) (96) to visualize structural variants greater than 300 kbp in length. This method of identifying large segregating CNVs is constrained in that it can only identify CNVrs that contain a presence variant on haplotype 1 of the reference. Since haplotype 1 is larger than haplotype 2, it is more likely to contain more presence variants. Secondly, to create a strong signature of divergence within the local PCA, a large number of individuals must contain the absence variant, meaning that presence variants will also be rare. Furthermore, closer inspection of segregating CNVrs within the reference indicated that some CNVrs may consist of complex SVs, containing inversions and translocations, which our identification method using WGS failed to identify.

Since cnv-chr4a was surprisingly large and segregating in our diploid reference, we conducted further analyses to understand its genomic makeup. Firstly, we analyzed TE content within this region. We identified TEs using EDTA (97) and used RepeatMasker v4.1.1 (98) to obtain a summary of various TE families within this region relative to the rest of chromosome 4. We then performed a synteny analysis to determine if the small number of genes in that region were collinear with other genomic regions and were therefore an ancestral arrangement, or if they were novel combinations of genes. We used McScanX v97e74f4 (99) to determine syntenic gene groups resulting from a self-alignment of protein sequences on haplotype 1 using blastp (-evalue 1e-10) in BLAST v2.7.1 [(100)]). We calculated read-depth on chromosome 4 for three aligned Ambrosia outgroup species (30). Calculating the mean read-depth exclusively within the cnv- chr4a region allowed us to ascertain the likely ancestral state of this structural variant.

### Temporal CNV changes

To analyze shifts in CNV frequency over both time and space, we used generalized linear models with the glm function in R (101) to predict presence/absence counts in populations from range (Europe or North America), sample year and sample latitude, including significant interactions between the three variables. PC1, which was calculated from a covariance matrix of 10,000 SNPs randomly sampled from outside of genes and putative inversions identified by Battlay et al. (15), was included in each model to control for population structure. Model significance was tested with the anova function using a type-3 test (car v3.1-2 package [(102)]). The models were reduced in a stepwise fashion, removing non-significant interactions until all remaining interactions (if any) were significant (p < 0.05). The emtrends function (emmeans v1.10.2 package [(103)]) was used to test directionality and obtain confidence intervals within interacting predictors.

### CNV-trait associations

To investigate associations between copy number variants and traits, we measured associations between 29 traits (in 121 samples for which trait data were available [(43)]) and the coverage windows described above. For each window, we fit linear models, using lm in R (101), between each sample’s normalized window depth and each trait value. To account for population structure, the first principal component of neutral covariance was added to the model. As previously, this was obtained by calculating a covariance matrix on 10,000 putatively neutral sites that were outside gene and inversion regions using PCangsd. A PCA was performed on this covariance matrix using prcomp in R (101). We assessed significance using a Bonferroni-corrected significance threshold: 0.05 divided by the number of windows tested (17,855). We performed CNVr-trait associations using EMMAX (v.beta-7Mar2010 [(104)]) to identify associations between the CNV regions, using a covariance matrix consisting of the same 10,000 putatively neutral SNPs in previous analyses in this study to correct for population structure.

### Gene Ontology analyses

Gene ontology enrichment analyses were performed using the R package topGO (105), using GO terms from A. thaliana TAIR 10 BLAST results. Fisher’s exact test, using a significance threshold of p < 0.05, was used to identify GO terms enriched within candidate gene lists relative to Q_ST_ outliers, as well as genes within CNVr.

## Acknowledgements

We thank Stephen Wright for his advice in the early stages of this project, and we thank Keyne Monro for her advice and expertise in developing the Q_ST_-F_ST_ analysis. We also thank Claire Mérot, an anonymous reviewer, and the editor for their thoughtful comments which greatly improved the manuscript.

## Data Accessibility

Sequences used in reference genome assembly and annotation are available from NCBI under BioProject ID PRJNA819156. The phased diploid genome assembly is available from NCBI under BioProject IDs PRJNA929657 and PRJNA929658. The haplotype 1 gene annotation GFF file is available from Figshare [https://doi.org/10.6084/m9.figshare. 19672710.v1]. Individual sample resequencing data are available from ENA under BioProject IDs PRJEB48563, PRJNA339123 and PRJEB34825.

## Code availability

All code for analyses performed in this work is available from Github [https://github.com/jonrobwil/ragweed_cnv]

